# Uncovering the genomic basis of infection through co-genomic sequencing of hosts and parasites

**DOI:** 10.1101/2022.12.05.519109

**Authors:** Eric Dexter, Peter D. Fields, Dieter Ebert

**Affiliations:** Department of Environmental Sciences, The University of Basel; Basel, 4051, Switzerland

**Keywords:** Coevolution, parasite, GWAS, daphnia, pasteuria, genome-to-genome, co-genomics

## Abstract

Understanding the genomic basis of infectious disease is fundamental objective in coevolutionary theory with relevance to healthcare, agriculture, and epidemiology. Models of host-parasite coevolution often assume that infection requires specific combinations of host and parasite genotypes. Coevolving host and parasite loci are therefor expected to show associations that reflects an underlying infection/resistance allele matrix, yet little evidence for such genome-to-genome interactions has been observed among natural populations. We conducted a study to search for this genomic signature across 258 linked host (*Daphnia magna)* and parasite (*Pasteuria ramosa)* genomes. Our results show a clear signal of genomic association between multiple epistatically-interacting loci in the host genome, and a family of genes encoding for collagen-like protein in the parasite genome. These findings are supported by laboratory-based infection trials, which show strong correspondence between phenotype and genotype at the identified loci. Our study provides clear genomic evidence of antagonistic coevolution among wild populations.

## INTRODUCTION

Parasites (including pathogens) infect nearly all forms of life, acting as a fundamental biological force that shapes the evolution of their hosts and drives the dynamics of entire communities (Poulin 2011; Schmid-Hempel 2021). The evolutionary processes among host and parasite are tightly interconnected: As a host population evolves in response to the threat caused by its parasites, so too does the parasite evolve in response to its host. This antagonistic coevolution may be responsible for much of the genetic diversity found in natural populations, promoting local adaptation and population differentiation (Morand 2015; Ebert and Fields 2020). Various models of host-parasite coevolution have been developed to accommodate the fact that phenotypic traits under selection may have different genomic architectures. Under highly specific models of host-parasite interactions, such as the matching-allele and the gene-for-gene models, infections are only possible between specific combinations of host and parasite genotypes (Agrawal and Lively 2002). Parasites will therefore experience high fitness when compatible alleles are common in the host population, and low fitness when such alleles are rare, thus driving a time-lagged cycle of negative-frequency dependent selection which is sometimes known as the Red Queen model of coevolution (Ebert and Fields 2020).

Red Queen evolutionary dynamics are predicted to yield several diagnostic characteristics in both host and parasite genomes. As negative-frequency dependent selection promotes the maintenance of genetic diversity, affected loci are expected to exhibit locally elevated nucleotide diversity relative to other genomic regions (Ebert and Fields 2020). This is because selection is tied to allele frequencies, with common variants being selected against. Highly specific models of host-parasite interactions also predict that certain combinations of host and parasite alleles are unlikely to be found in infected hosts, even if both alleles occur in the population at large. Coevolving host and parasite loci are therefore expected to show a statistical pattern of association that reflects the underlying infection/resistance allele matrix (Ebert and Fields 2020). The non-random association of alleles across the genomes of two interacting species can be considered as a form of interspecies linkage disequilibrium, more succinctly “interlinkage.” Interlinkage is a predicted consequence of genomic interactions between hosts and parasites and thus direct evidence for ongoing coevolutionary interactions of the antagonists. Despite arising as a fundamental prediction of host-parasite coevolution, there is little empirical evidence of interlinkage among naturally coevolving populations. Here we present clear evidence of interlinkage among natural populations of the planktonic crustacean *Daphnia magna* and its endoparasitic bacterium *Pasteuria ramosa* (Firmicutes).

*Daphnia magna* is a common planktonic microcrustacean that inhabits standing fresh and brackish waters across most of the northern hemisphere. Wild populations of *D. magna* are often heavily parasitized by a wide variety of bacteria, virus, and microsporidian species, including the endospore-forming bacteria *Pasteuria ramosa* (Ebert 2005). *Daphnia magna* ingest planktonic spores of *P. ramosa* during filter feeding, which enter the digestive tract along with food and other particulate matter and may attach to the gut lining. After attachment and penetration of the host cuticle, *P. ramosa* rapidly proliferate in the body cavity of the host (Ebert et al. 1996). During *P. ramosa* infections, the normally transparent host body cavity is completely filled with opaque spores which will be released into the environment upon death of the host. *P. ramosa* infection spreads exclusively through horizontal transmission from host to host.

The *D. magna* / *P. ramosa* infection process appears to be governed by highly specific interactions, such that *D. magna* genotypes (aka. clones) are either completely susceptible or completely resistant to particular strains of *P. ramosa* (Bento et al. 2017; Bento et al. 2020). This binary outcome is strongly linked to the attachment of the parasite to the host cuticle. This highly-specific attachment step is hypothesized to be governed by interactions between *P. ramosa* surface proteins and highly-polymorphic extracellular proteins at the target tissue of the host (Andras et al. 2020). Extensive attachment testing with a wide range of host and parasite isolates has shown that susceptibility is almost perfectly correlated with attachment, with little to no variation due to environmental or host condition (Duneau et al. 2011).

Previous studies have shown that *D. magna* resistance to *P. ramosa* attachment is governed by multiple epistatically interacting loci distributed across the *D. magna* genome. Genetic crosses of *D. magna* clones with differing *P. ramosa* resistance phenotypes have thus far identified six resistance loci: the A, B, C, D, E, and F loci. The A, B, and C loci cluster tightly together on chromosome 4 to form the ABC supergene, which shows major structural polymorphism between haplotypes, including long stretches of nonhomologous sequences that appear to be non-recombining (Bento et al. 2017). The F locus is closely located to the ABC cluster, while the D and the E loci are distributed elsewhere across the *D. magna* genome, but epistatic interactions between combinations of these loci have been described (Bento et al. 2017; Ameline et al. 2021).

The loci which govern infection in the *P. ramosa* genome are less well understood, but there are indications that a highly expanded family of genes encoding collagen-like proteins may be a determinant of infection outcome. Collagen-like proteins are known to be strongly tied to the infection process in a number of bacteria species by mediating attachment to host cells (Qiu et al. 2021). Pasteuria collagen-like (PCL) genes are widely distributed across the *P. ramosa* genome in triplet clusters, potentially the consequence of repeated duplications of a highly mobile transposable element (McElroy et al. 2011). A previous study found that variation at one of these PCLs correlated perfectly with the ability of certain *P. ramosa* isolates to infect a specific *D. magna* genotype (Andras et al. 2020). Given that *P. ramosa* spores are coated with a thick layer of fibers hypothesized by be composed of collagen-like proteins (Andras et al. 2020), and that spore attachment to any of several sites on the host cuticle appears to be the primary determinant of infection outcome (Duneau et al. 2011; Fredericksen et al. 2021), these PCL genes are promising candidates for sites of interlinkage with host resistance genes.

Previous studies of the *D. magna* / *P. ramosa* system have examined loci governing the infection process through experimental contexts which allow for genomic variation only for the host or for the parasite. Although multiple resistance genes in the host and one infectivity gene in the parasite are known, the manner in which these genes interact with each other is not yet understood. Such interactions are at the core of matching-allele and gene-for-gene models, thus forming the most fundamental aspect of coevolutionary dynamics. A number of different statistical approaches have been developed to detect signals of coevolution, both at the genomic level, and the phylogenetic level (Legendre et al. 2002; Carlson et al. 2008; Wang et al. 2018; Märkle et al. 2021). The co-GWAS approach is an extension of single-phenotype GWAS (Genome Wide Association Study), which simultaneously considers genetic information from host and parasite. Through the combination of clinical data and genome-scale SNP arrays, co-GWAS has been used to identify genomic variants of clinical significance for HIV infection (Bartha et al. 2013), Hepatitis C (Ansari et al. 2017), and pneumococcal meningitis (Lees et al. 2019). In order to identify potentially interlinked loci among our system, we conducted a co-genome wide association study (co-GWAS) using wild collected *D. magna* and their naturally associated *P. ramosa* parasites.

## RESULTS

### Sequencing and mapping

We sequenced 258 *D. magna* naturally infected with *P. ramosa* collected during an epidemic across one season from a single pond, and separated the resulting reads by mapping against reference genomes for the host and parasite. Overall rates of contamination were low for whole-organism sequencing, with 92.8 % (11.0 SD) of reads mapping to either the host or parasite genome (S1 Fig). Mean depth of coverage was 31.0 (SD = 13.0) for *D. magna* and 318 (SD = 217) for *P. ramosa*. This difference in mean coverage reflects the smaller genome size of the parasite, which is approximately 2 Mb versus the approximately 230 Mb host genome. Ten of the samples showed low coverage for either the host or parasite genome and were excluded from further analysis (S1 Fig).

### Genomic variation

We identified 21,933 variant positions (SNPs, indels, and SNP/indel combinations) in the *P. ramosa* sequence data. These variants were filtered for base and mapping quality, excessive read depth, and bi-allelic status, resulting in 10,310 variant positions retained for downstream analysis. For the much larger *D. magna* genome, an initial call set of 4,142,757 variants was reduced to 2,211,993 after identical filtering parameters were applied. Population-wide nucleotide diversity (π) was 0.26 for *D. magna* with per sample missingness rates < 1 %. Estimates of genetic diversity within *P. ramosa* samples varied dramatically, likely reflecting that some hosts had been infected by multiple *P. ramosa* genotypes, while others were subject to single infections (Fig 1A). Eight samples were monomorphic at nearly all sites, thus indicating the presence of only a single haploid *P. ramosa* genotype. The remaining samples showed variable rates of *P. ramosa* polymorphism with extreme cases showing variation at up to 50 % of segregating sites, suggesting multiple infections by dissimilar genotypes (Fig 1A).

**Figure 1.**
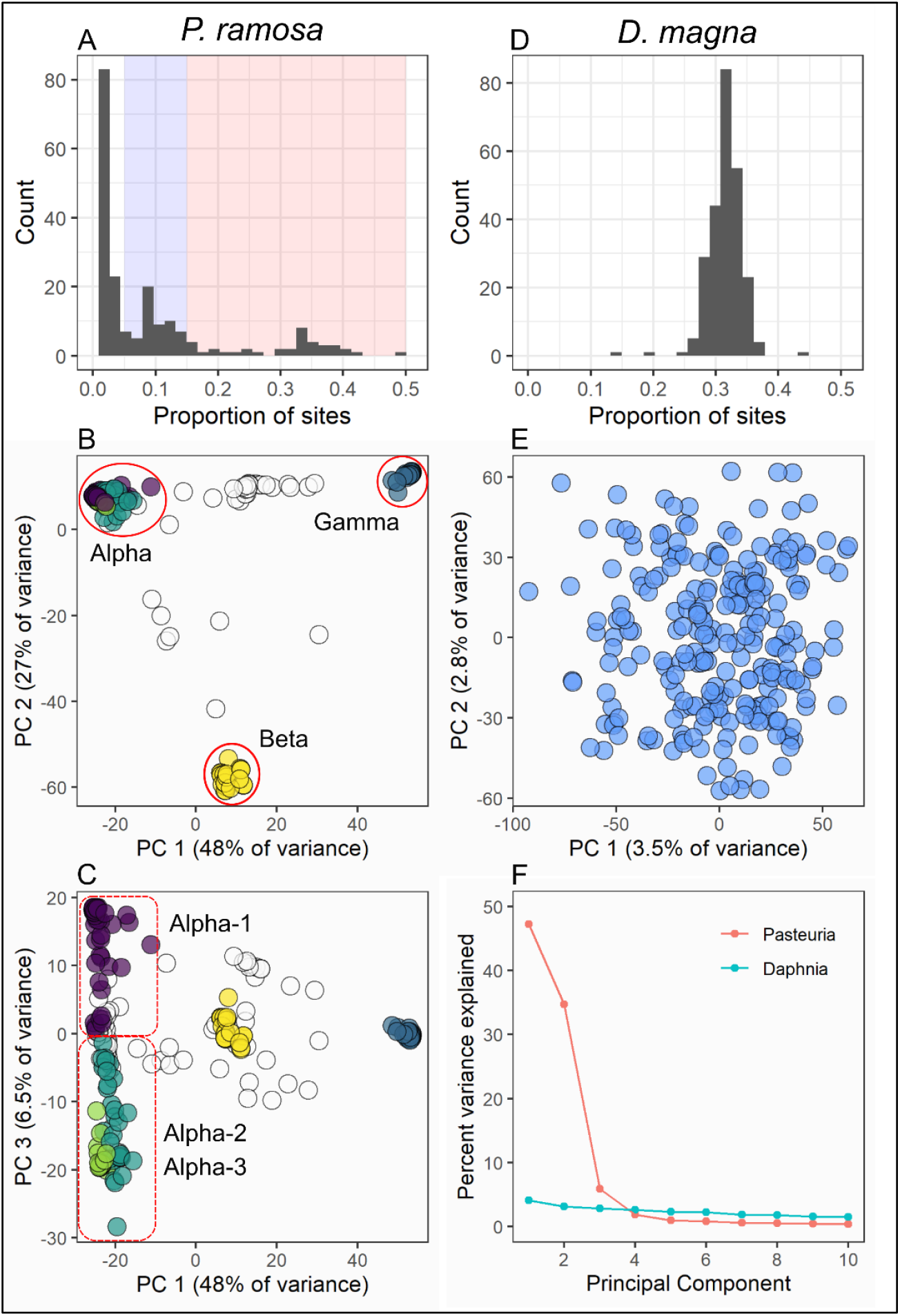
**G**enomic diversity and population structure of samples used in this study. **(A)** Per sample levels of genomic diversity for *P. ramosa*, expressed as the proportion of population-wide variant sites showing within sample polymorphism. Single genotype infections of the haploid *P. ramosa* are expected to show values near zero, while larger values indicate multiple infections. Values > 0.15 represent infections from multiple distinct pasteuria lineages (shaded in red), with lesser values indicating multiple infections of closely related genotypes (blue). **(B)** PCA ordination of parasite genomes (N = 248) recovered from infected hosts. Parasites cluster into three primary genomic groups (*Alpha, Beta, Gamma*), with membership derived by DAPC clustering of whole-genome data. The colors of the points represent DAPC clusters. Open circles represent multiple infections containing mixtures of genotypes from different clusters (red shaded area on panel A). **(C)** The same PCA ordination rotated to show PCs 1 and 3. The *Alpha* cluster shows relatively continuous variation across the PC 3, which DAPC clustering hierarchically sub-divides into *Alpha-1* and *Alpha-2* / *Alpha-3*. Note the relatively modest contribution to total genomic variation by PC 3 relative to the first two PCs. **(D)** Per sample levels of genomic diversity for *D. magna*, where each sample represents a single diploid individual. **(E)** PCA ordination of *D. magna* samples (N = 248) plotted across the first two principal components. Note the low percentage of variance explained by the primary axes and the lack of overall structure. DAPC clustering for *D. magna* best supports a single cluster. **(F)** PCA scree plots for *P. ramosa* and *D. magna*.

### Population structure

We examined host and parasite samples for hidden population structure using principal component analysis (PCA) and Discriminant Analysis of Principle Components (DAPC). The *D. magna* samples showed no discernable population structure, matching our expectations for a sexually reproducing population contained within a small pond. The strongest principal component for *D. magna* explained 3.5 % of total genomic variation, with subsequent PCs explaining incrementally smaller values (Fig 1E, F). K-means clustering based on genetic distance showed the strongest support for a single *D. magna* cluster, which is readily observable upon visual inspection of the PCA ordination (Fig 1E).

In contrast to the host, the majority of genomic variance for *P. ramosa* could be summarized by the first two principal components, which together explained 77 % of total variance (Fig 1B). Visual inspection of the first two axes of the PCA ordination showed the *P. ramosa* samples arranged as a triangular cloud with three distinct genomic clusters in the corners, which we have labeled as *Alpha, Beta*, and *Gamma* (Fig 1B). Samples infected by multiple *P. ramosa* genotypes (Fig 1A) were located between the clusters, primarily along the edges of the triangle. K-means clustering based on genetic distance supported the division of single-infection samples into the *Alpha, Beta, Gamma* clustering scheme, with the possibility for further subdivision of the *Alpha* lineage into three sub-clusters (Fig 1B, 1C). Admixture analysis of our *P. ramosa* samples with K=3 groups showed identical placement of samples respective to the *Alpha, Beta*, and *Gamma* clustering scheme as derived from DAPC analysis (S2 Fig). For the sake of clarity, we hereafter refer to the three *P. ramosa* genomic clusters as “lineages” without reference to specific taxonomic rank.

The examination of further parasite principal components revealed a small degree of relatively continuous within-lineage variation, potentially reflecting the process of recombination within lineages (Fig 1C, S2). Recombination is not well understood in *P. ramosa*, but we could observe its signature in the form of linkage disequilibrium dissolution with distance across the *P. ramosa* genome (S3 Fig). Admixture analysis indicated strong support for recombination within lineages, but much less between the three primary lineages (S2 Fig). Therefore, the boundaries between sub-clusters should be understood as approximation rather than the clear divisions that we observe between the discrete *Alpha, Beta*, and *Gamma* lineages.

Population structure is an important consideration in GWAS modeling, and is often included as a covariate in the form of principal components or a relatedness matrix (Zhou and Stephens 2012; Chang et al. 2015). This paradigm can be extended to co-GWAS models through inclusion of principal components from both the host and parasite (Ansari et al. 2017; Naret et al. 2018). However, several unique features of our system require careful consideration regarding the treatment of population structure in a co-GWAS context. In this study, the *D. magna* samples originated from a single population absent of geographic subdivision (a small pond). Under this demographic scenario, the major principal components are likely to represent local genomic features rather than genome-wide structure. The inclusion of these PCs as covariates in co-GWAS model might therefore mask highly differentiated loci which are potentially of biological interest from model results. This is even more the case for the parasite, as the strong lineage structure that we found for *P. ramosa* may be tightly linked to (or even derive from) the infection phenotype of the parasite. If so, the top principal components for the parasite would represent a large portion of the biologically relevant aspects of its interaction with the host. Correcting for population structure might therefore mask the biological associations at the core of the interaction.

### Co-GWAS

In consideration of these aspects, we have produced three hierarchically nested co-GWAS models that examine host-parasite interlinkage under different assumptions regarding population structure, dominance, and host-specificity. Model 1 assumes that potentially cofounding demographic structure exists for both host and parasite, and thus the model contains principal components from both as covariates. Model 2 assumes that parasite population structure is a consequence of host-specificity, and thus the model contains principal components for only the host. Model 3 assumes that the host population is unstructured and omits all principal components, which greatly reduces the total number of parameters estimated. Many other biologically reasonable model structures could also be envisioned, but we found that these three models provided a good overall representation of the parameter space. Potential effects of host allele dominance could be tested only in model 3, as model structures which included further covariates experienced convergence problems due to model complexity. Given the large number of pairwise associations that are tested under each model (>10^9^) we have avoided setting a specific p-value threshold to assign statistical significance to any particular model results. Rather, the strongest peaks from each model were considered as candidate loci for phenotypic confirmation through laboratory-based infection trials.

Our first co-GWAS model contained controls for host and parasite population structure, with the top 10 principal components from the host (22 % total variance) and top 2 principal components from the parasite (82 % total variance) as model covariates. The results from this model showed a single strong peak in the daphnia genome located precisely at the location of a well-characterized locus for *P. ramosa* resistance - the ABC locus (Table 1). The ABC locus is considered a supergene (Bento et al. 2017), but we likely observe only the effect of the C locus here because the A locus shows no variation in our study population, and the B locus is nearly fixed for one allele (Ameline et al. 2021). Note that the A and B loci are not globally fixed and show greater polymorphism in some populations (Luijckx et al. 2014; Metzger et al. 2016; Bourgeois et al. 2021). This genomic region showed a strong signal of interlinkage with a tightly clustered set of genes encoding triplets of collagen-like proteins in the *P. ramosa* genome (Fig 2). This small region contains the densest concentration of collagen-like genes in the *P. ramosa* genome, containing PCL triplets [9-10-11], [12-13-14], and [36-37-38] (Fig 2, S4). A single, apparently intragenic, SNP on the right arm of chromosome 5 of the host genome also appeared among the top results for this model (Fig 2, Table 1), but the complete absence of neighboring variants with similar small p-values indicates that this peak is relatively weak candidate (S5 Fig).

**Table 1.** Summary of top model results from each co-GWAS model. Rather than set a specific p-value thershold for statistical significance, the peaks described below represent the top 500 associations for each model as plotted in figure 2.

| Model | Peak position (*D. magna*) | Peak description (*D. magna*) | Peak position (*P. ramosa*) | Peak description (*P. ramosa*) |
| --- | --- | --- | --- | --- |
| 1 | Chromosome 4 (right arm)<br>Contig 11: 2084705 – 2382802 | Overlaps the C locus | 1565875 – 1600196 | Overlaps PCL triplets [9-10-11], [12-13-14], and [36-37-38] |
|  | Chromosome 5 (right arm)<br>Contig 71: 160694 | Single SNP, intragenic,<br>possible mapping error | 758063 | Aminofutalosine synthase (electron transport) |
| 2 | Chromosome 4 (right arm)<br>Contig 11: 2084705 – 2357097 | Overlaps the C locus | 1565739 – 1628258 | Overlaps PCL triplets [9-10-11], [12-13-14], and [36-37-38] |
|  | Chromosome 7 (right arm)<br>Contig 18: 1785587 – 1803392 | Overlaps the D locus | 318169 – 343599 | Overlaps PCL triplet [21-27-28] |
|  | Contig 32: 849103 | Single SNP, intragenic,<br>possible mapping error | 1347536 – 1382775 | IMP dehydrogenase and DNA topoisomerase subunit B |
| 3 | Chromosome 5 (left arm)<br>Contig 1: 6277551 – 6506684<br>Contig 67: 230263 – 233072 | Overlaps the E locus | 1588560 – 1588747 | Overlaps PCL triplet [36-37-38] |
|  |  |  | 4530 – 4707 | 3 Kb from PCL triplet [6-7-8] |
|  | Chromosome 5 (right arm)<br>Contig 25: 43625 – 1636178<br>Contig 53: 182869 – 443318 | Shows strong correlation with<br>resistance to *P. ramosa*<br>isolates P20 | 4530 – 8506 | Overlaps PCL triplet [6-7-8] |
|  |  |  | 319494 – 319584 | Overlaps PCL triplet [21-27-28] |
|  |  |  | 1519089 – 1520569 | Overlaps PCL triplet [24-25-26] |
|  |  |  | 1588269 – 1589748 | Overlaps PCL triplet [36-37-38] |
|  | Chromosome 7 (left arm)<br>Contig 3: 4270730 – 4354576 | Mostly uncharacterized genes | 1181208 – 1205486 | Overlaps PCL triplet [15-16-17] |
|  | Chromosome 7 (right arm)<br>Contig 18: 1785587 – 1802479 | Overlaps the D locus | 1516194 | 1 Kb distant from PCL triplet [24-25-26] |

**Figure 2.**
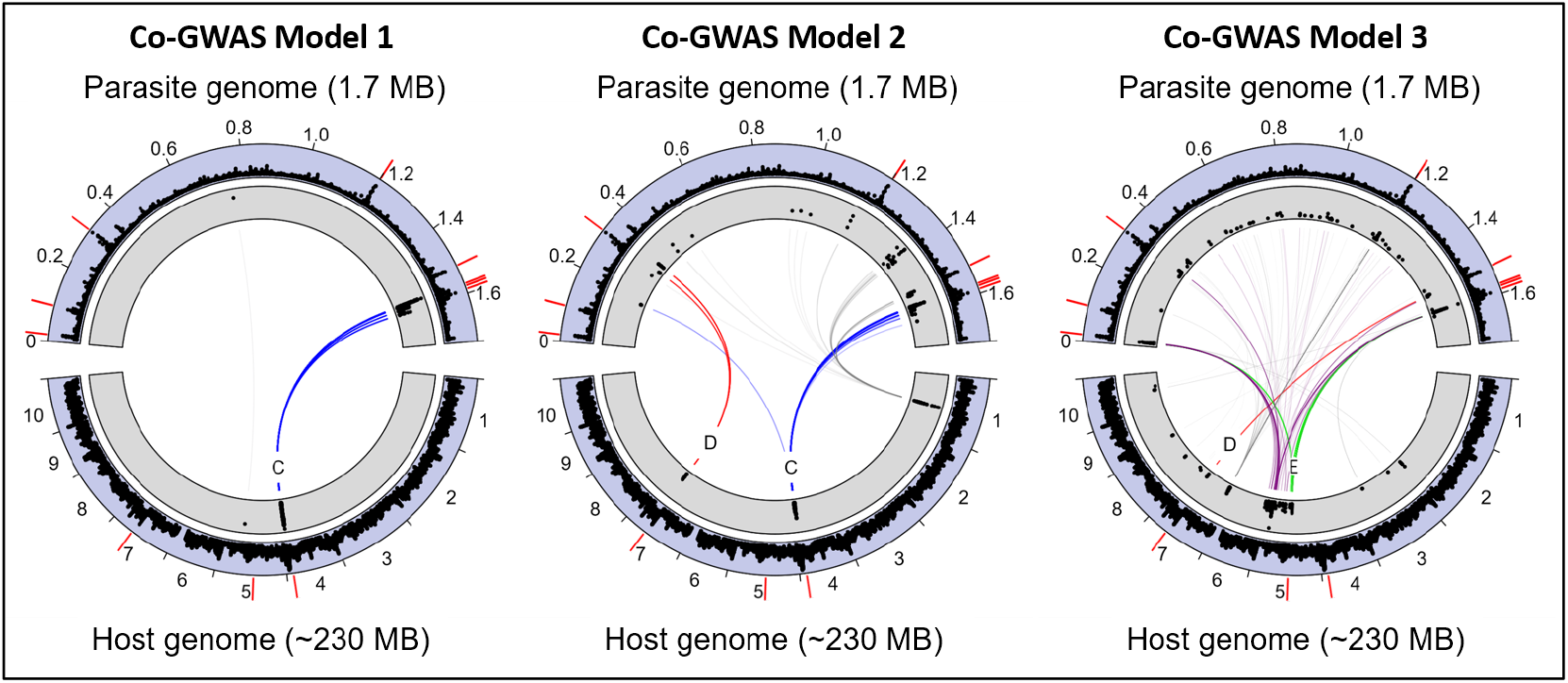
The 500 strongest associations for each co-GWAS model between variants in the host genome (bottom ring segments) and the parasite genome (top segments). Inner ring segments (gray background) indicate the scaled genomic position (x-axis) and strength (y-axis) of the association, expressed as the p-value of the co-GWAS score on a -log_10_ scale. Outer ring segments (with blue background) indicate nucleotide diversity (π), scaled from 0 to 1. Darker pathways indicate a greater number of associations, with known resistance loci of the host being shown in blue, red, and green (C, D, and E locus, respectively). Outer red bars indicate the position of PCL gene triplets in the *P. ramosa* genome and resistance genes in the *D. magna* genome; the outer ring of the host genome displays chromosome numbers. Model 1 shows a strong association between the host C locus and a dense cluster of PCL gene triplets in the parasite genome. Model 2 shows the C locus (as in model 1), but also shows a strong association between the host D locus and a different PCL gene triplet in the parasite genome. In model 3 the strongest associations occur on two regions of chromosome 5: One region coincides with the previously described E locus for pasteuria resistance, the other corresponds to a previously undescribed candidate locus for pasteuria resistance which may interact with the E locus (see main text). Lines shown in purple pertain to a new candidate locus for *Pasteuria* resistance.

Our second co-GWAS model included covariates from the host only, thereby allowing parasite populations structure to be incorporated into the genomic terms of the model. This model showed the same C locus / PCL interlinkage observed in model 1, and a further strong signal of interlinkage precisely at the location of the D locus for *P. ramosa* resistance (Fig 2). The D locus appears to be tightly interlinked with a narrow genomic region containing the PCL gene triplet [21-27-28] (Table 1). We also observed a single, apparently intragenic, host SNP on chromosome 1 among the top results for this model. Like the isolated SNP observed in model 1, the complete absence of neighboring variants with similar p-values suggest that this peak is relatively weak candidate.

Our third co-GWAS model accounted for the fact that our host population is best represented as a single well-mixed population, and further omitted all principal components for the host. Model 3 also allowed for dominance effects in the host genotype, which could not be incorporated in previous models due to convergence problems with the larger number of parameters estimated. The strongest signatures of interlinkage in model 3 were again located between previously-characterized pasteuria-resistance loci in the host genome and PCL gene triplets in the parasite genome (Fig 2). The E locus region for *P. ramosa* resistance appeared as one of the largest peaks in model 3, and showed strong interlinkage with PCL triplets [6-7-8], and [36-37-38]. We also observed a strong signal of interlinkage between the D locus for *P. ramosa* resistance and PCL [24-25-26]. The C and D loci showed clear associations with PCL gene triplets as observed in the other co-GWAS models, but with a relatively lower magnitude than other peaks in model 3 (S6 Fig).

In addition to these known pasteuria-resistance loci, we observe a strong signature of interlinkage in model 3 at two previously-uncharacterized positions in the host genome. The largest of these peaks was found across a broad section of the right arm of chromosome 5, and was associated with PCL triplets [6-7-8], [21-27-28], [24-25-26], and [36-37-38]. The extreme width of this peak is unusual, and perhaps indicates some type of structural polymorphism. The second uncharacterized peak in model 3 occurred between a region on the left arm of host chromosome 7 and PCL triplet [15-16-17]. Neither of these loci have been previously observed to relate to *P. ramosa* resistance in laboratory experiments, but they appear here as strong candidates for further investigation based on the width of the peaks and their highly specific PCL triplet association.

### Phenotypic confirmation

In order to assess the correspondence between host and parasite phenotype, we conducted a series of infection trials using clone lines established from our wild-collected daphnia and a panel of *P. ramosa* isolates. This panel was comprised of five laboratory-cultured *P. ramosa* isolates (C1, C19, P15, P20, and P21), which in various combinations reveal the phenotype of the known pasteuria-resistance loci (Luijckx et al. 2013; Bento et al. 2017; Bento et al. 2020; Ameline et al. 2021). For example, the allelic state at the *D. magna* C locus is defined in relation to susceptibility to *P. ramosa* isolates C1 and C19, with resistance being dominant (Bento et al. 2017). Three resistotypes comprised 81% our sequenced samples: [RRSRS] (52%), [SSSSS] (21%), and [RRSSS] (8%). These values match expectations from previous studies of the *P. ramosa* epidemic at lake Aegelsee which observed the same three resistotypes in roughly similar proportions (Ameline et al. 2021). Twelve additional rare resistotypes were observed with most present in only a single individual. With a much larger sample size Ameline et al. (2021) also observed that certain multi-locus resistotypes are rarely or never observed. We observed variation in respect to resistance/susceptibility for all tested isolates, although most daphnia (93%) were susceptible to isolate P15. Only one specimen showed resistance to all five laboratory isolates. It is interesting to note that the most common resistotypes were not necessarily those which conferred the broadest pattern of resistance.

We then applied DAPC clustering to the genomic regions corresponding to each of the resistance loci, and grouped the genomic clusters by the relevant resistotype to define functional genotype groups (S7, S8, S9 Tables). Up to 10 genotype clusters could be detected at each resistance locus, likely representing several haplotypes in various combinations with each other. Our goal was to identify the minimum number of clusters needed to differentiate resistance phenotypes, and in all cases, we found that three clusters gave us the best segregation of resistotypes. These three clusters could then be collapsed into two groups representing primarily resistant or primarily susceptible phenotypes. This classification scheme of three genotypes clusters segregating into two phenotypes likely reflects underlying dominance effects in heterozygous loci as predicted by our genetic models, but the presence of multiple haplotypes and uncertainty about the precise nature of the variants at these resistance loci limits our inference. Among our samples we observed perfect correspondence between DAPC genotype cluster at the C locus and resistance phenotype, with one of genotypes (C’) showing complete susceptibility to C1 and C19 and the other (C) showing resistance to C1 and C19 (S6 Table). We observed only partial correspondence between resistotype and genotype for the D and E loci. In both cases, one genotype cluster (D or E) corresponded to exactly one resistance phenotype, but the alternative genotype cluster (D’ or E’) corresponded to multiple resistance phenotypes (S7, S8 Table). This imperfect correspondence between genotype and phenotype does not appear to be an artifact of our clustering methodology or dominance effects, as apparently identical genotypes were associated with multiple resistance phenotypes.

### Uncharacterized loci recovered from co-GWAS

Our co-GWAS models recovered a strong signal of interlinkage at two previously uncharacterized loci in the *D. magna* genome. The strongest of these peaks spanned a large (∼ 4 Mb) region on the right arm of chromosome 5, while the other spanned ∼ 100 Kb on the left arm of chromosome 7. Given the imperfect sorting of D and E locus genotypes into their expected phenotypes, we hypothesized that one or both of these uncharacterized loci could represent an unknown pasteuria-resistance locus that epistatically interacts with either locus. We therefore performed a series of simple GWAS experiments to determine if these loci correspond to different pasteuria resistance phenotypes. We discovered that the candidate locus on the right arm of chromosome 5 is strongly associated with the resistotype RRS relative to the *P. ramosa* isolates C1, C19, and P20 (Fig 3). This is the canonical resistotype used to identify the allelic state at the E locus in laboratory-based experiments (Ameline et al. 2021), and provides further evidence that this previously uncharacterized locus may epistatically interact with other loci to affect resistance phenotype. We could not identify a similar correlation between resistance phenotype and the second uncharacterized locus on chromosome 7, but future experiments are planned to further examine this possibility through an expanded panel of *P. ramosa* isolates.

**Figure 3.**
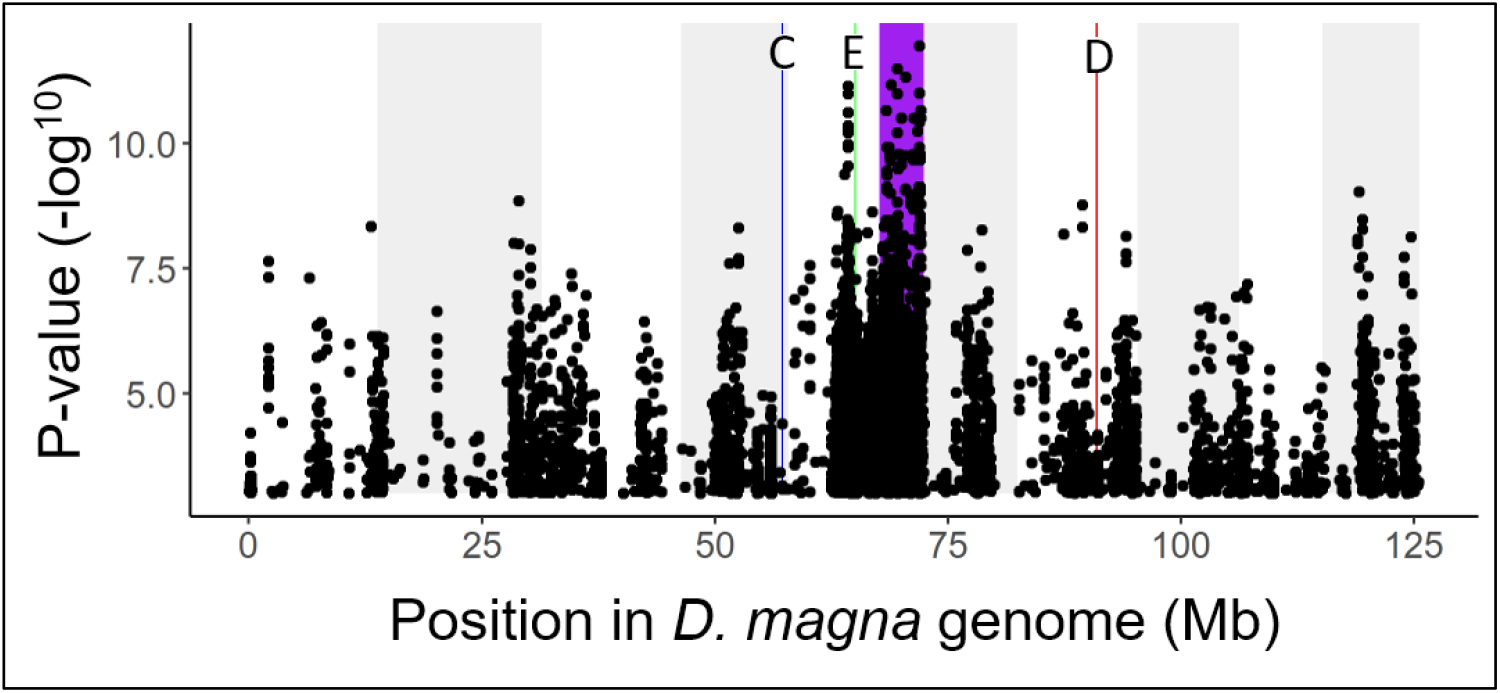
Single phenotype GWAS for resistance to the *P. ramosa* isolate P20. Resistance or susceptibility to this isolate is diagnostic of the allelic state at the E locus. The positions of the C locus (blue), D locus (red), E locus (green) and new candidate locus for *Pasteuria* resistance (purple) are indicated by vertical bars. Our results show a strong peak at the position of the E locus (green) as expected from previous studies. However, we observe a much stronger peak on the right arm of chromosome 5 (purple) that coincides with the primary peak from our third co-GWAS model. This result provides further evidence of a uncharacterized pasteuria resistance locus at this position, which may epistatically interact with the E locus.

### Genotype – Genotype matrix

Host specificity in the *D. magna* / *P. ramosa* system is hypothesized to arise from relatively simple host-parasite infection matrices, such as a matching allele matrix, but the presence of multiple interacting loci in both host and parasite genomes has the potential to yield considerable complexity. Nonetheless, our functional grouping of hosts according to resistance locus genotypes revealed strong associations with the parasite genomic clusters. Chi-squared tests of genotype contingency tables showed significant associations for the C locus (Χ^2^ = 16.4, p = 2.5^-3^), D locus (Χ^2^ = 32.5, p-value = 1.5^-6^), and E locus (Χ^2^ = 55.2, p = 3.0^-11^), with deviations from expected values resembling a matching allele model (Fig 4). For example, we observed that *D. magna* carrying the D genotype were most likely to be infected by parasites from the *Gamma* lineage, but apparently immune to infection by *Gamma* when carrying the D’ genotype (Fig 4). Similarly, the E genotype appears to confer strong resistance to the *Beta* lineage, while the alternative E’ genotype confers strong susceptibility. This specificity remains observable at finer scales, with various sub-clusters of the *Alpha* lineage showing strong differences in ability to infect different C and D locus genotypes.

**Figure 4.**
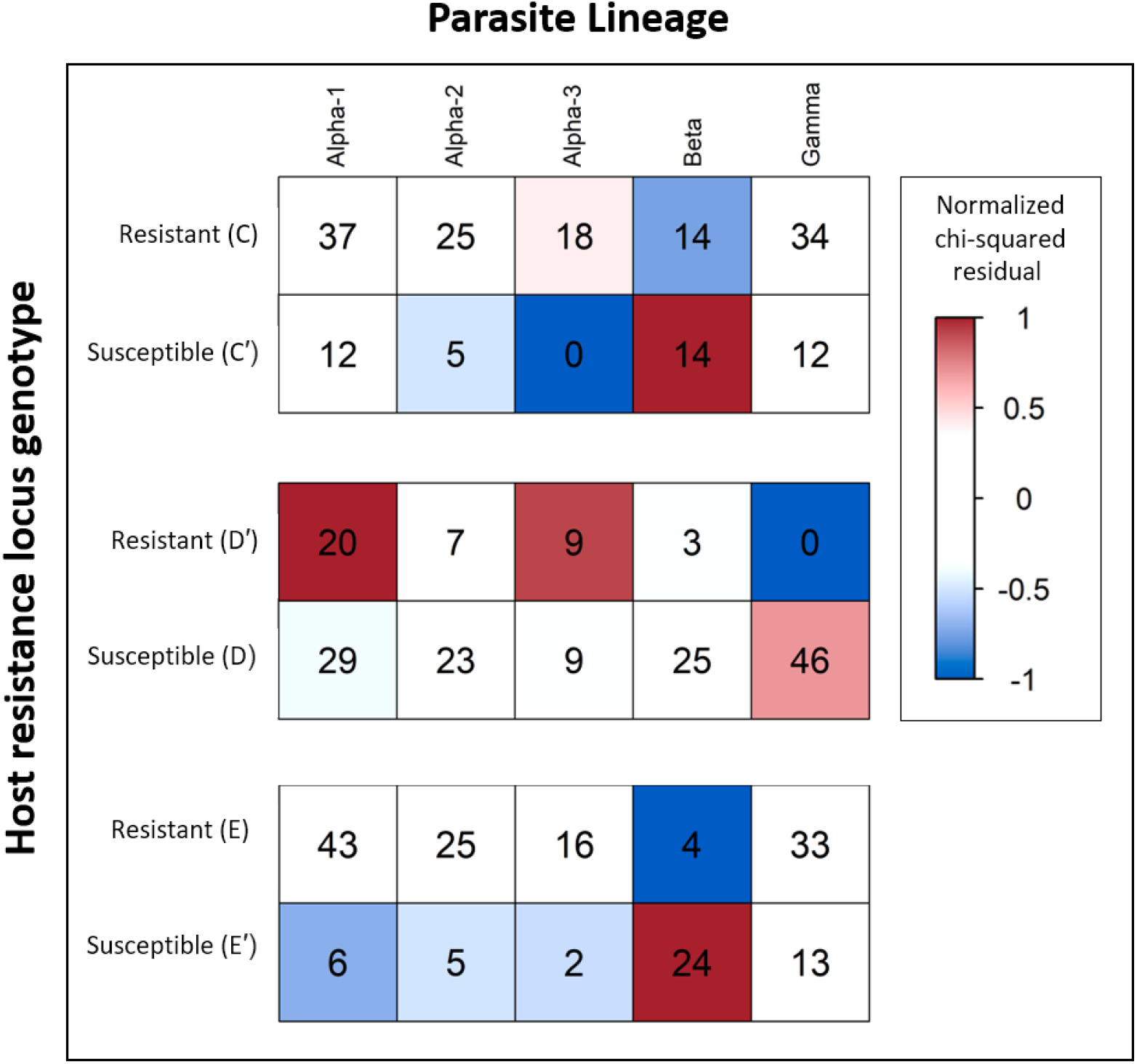
Contingency tables of genotype clusters at host resistance loci (derived via DAPC) versus *P. ramosa* genomic cluster. Cells are colored according to the magnitude and direction of deviation from the expected number of observations (Pearson residuals), with host and parasite combinations occurring more often than expected shown in red, and less often in blue. Host genotype clusters are characterized as resistant or susceptible relative to their diagnostic *P. ramosa* isolates – none are universally susceptible or resistant. Residuals for host-parasite combinations were calculated from a chi-squared contingency table test (p < 0.001 for all matrices, see text for full result) and then rescaled from -1 to 1 for color plotting. Numbers inside cells are the counts for each combination. Multiple infections (open circles on panel B) are not included here, resulting in a total count of 171 in each table.

## DISCUSSION

### Summary

Our results showed a clear signal of genomic interlinkage between several loci in the *D. magna* genome and a set of genes encoding for collagen-like protein in *P. ramosa*. Our findings were supported by our *P. ramosa* attachment assays, which showed a strong correspondence between phenotype and genotype at the recovered resistance loci. The fact that our co-GWAS model recovered all known resistance loci of daphnia that segregate in our study population stands as a strong biological validation of our results, as experiments describing these loci employed genetic crosses and phenotyping experiments using a laboratory-based panel of hosts and parasites (Metzger et al. 2016; Bento et al. 2017; Bento et al. 2020; Ameline et al. 2021). Our co-GWAS models recovered these loci using only genomic data from wild-collected specimens, with phenotypic data employed only for post-hoc confirmation. While most of these loci in the host were known previously, their interactions with infectivity loci in the parasite were entirely unknown. Our experiments revealed that these are not general resistance and infectivity loci, but rather their effect is contingent on the allelic state at interacting loci in the host or parasite. We also identified a new candidate locus for pasteuria-resistance in the host genome which strongly correlates with resistance to *P. ramosa* isolate P20. A further important finding is that collagen-like gene triplets in the *P. ramosa* genome (but not single collagen-like genes) are key determinants of infectivity. These new candidate loci for infectivity and resistance will be examined through future experiments with genetic crosses and QTL mapping.

### Co-GWAS model differences

Our co-GWAS modeling revealed that different assumptions regarding population structure, dominance, sample relatedness, and other factors yielded important differences in the results. Therefore, we present not a single model that shows only a snapshot of this parameter space, but present three co-GWAS models which covers the range of variation that we observe under a range of biologically reasonable assumptions. Our first co-GWAS model includes principal components from both the host and parasite as covariates. As parasite lineage structure is strongly correlated with the first two principal components of the *P. ramosa* genome, we primarily observed the effects of within-lineage variation in model 1. This is supported by our host genotype – parasite lineage association tables (Fig 4) which show that the effect of the host’s C locus is most apparent across sub-groups within the *Alpha* lineage.

Our second co-GWAS model includes covariates from the host only, thereby allowing parasite populations structure to be incorporated into the genomic terms of the model. This approach could be considered an analytical mistake, as it could increase the rate of false positives due to non-adaptive demographic effects such as co-dispersal of host and parasite (Naret et al. 2018). However, under such a scenario all lineage specific alleles would show an equally strong association. We do not observe this phenomenon in our results for model 2, rather we observe clearly distinct peaks at specific regions of the host and parasite genome which contain genes previously hypothesized to govern the infection process. Model 2 reflects the hypothesis that population structure in *P. ramosa* is a direct result of genetic interactions with its host rather than a confounding factor.

Our third co-GWAS omitted host and parasite population structure under the assumption that specific combinations of host and parasite genomes occur due to biological rather than demographic reasons. This assumption is reasonable given that all samples were collected from one population during one season, but would be advised against when populations are comprised of discrete demes or drawn from a wide geographic range. Model 3 also allowed for dominance effects in the host genotype, which could not be incorporated in the models 1 and 2 due to convergence issues arising from the larger number of parameters. In this model we observed strong peaks of interlinkage between pasteuria-resistance loci and PCL gene triplets, and identified two additional candidate loci in the host genome showing similar signals of interlinkage. Subsequent laboratory experiments showed one of these loci to be highly correlated with resistance to *P. ramosa* isolate P20.

Although each of these models vary to some degree in specific details, we observe a consistent pattern of pasteuria-resistance genes in the host showing a highly specific association with collagen-like genes in the *P. ramosa* genome that are arranged in a conserved triplet configuration. The major differences between these models can be readily interpreted, and to some degree, predicted by the biology of the system. As the E locus loads heavily on the top principal components of *D. magna* in our study population (S11 Fig), we expect that the effect of this locus should detectable only in model 3. Similarly, model 1 is constrained in such a manner that only within-lineage associations should be observable, which is exactly what the model shows. The C and D loci remain detectable in model 3 (S6, S12 Fig), but show a much weaker signal than the E locus, reflecting previous observations that the E locus appears to be the primary determinant of pasteuria resistance in our study population (Ameline et al. 2021; Ameline et al. 2022).

### Discovery of P. ramosa lineages

Our discovery of three distinct evolutionary lineages of *P. ramosa* within the study site was a particularly unexpected result of this project. Although most daphnia hosts were infected by only a single lineage of pasteuria, simultaneous infection by multiple lineages of pasteuria were observed. Given that we detected strong evidence for recombination within our study population (S2, S3 Fig), a presently unknown mechanism is clearly maintaining some form of reproductive isolation between these lineages. The nature of this reproductive isolation between pasteuria lineages is an important question for further investigation. We intend to conduct broader surveys of pasteuria populations to determine if additional lineages occur globally. The discovery that *P. ramosa* isolates arise from discrete evolutionary lineages also allows us to assess how well our laboratory panel represents the diversity of genotypes present at our study site. Preliminary evidence with newly developed genetic markers indicates that the isolates C1, C19, and P20 belong to the *Beta* lineage, the isolates P15 and P21 belong to the *Gamma* lineage, and that the *Alpha* lineage is not represented in our standard panel. We are in the process of developing several new pasteuria cultures from the *Alpha* lineage to deploy in future studies, and a panel of genetic markers to differentiate lineages and potential sub-lineages.

### Signatures of balancing selection at interlinked loci

Negative-frequency dependent selection is a form of balancing selection that is expected to yield a genomic signature of locally elevated nucleotide diversity at the sites under selection (Ebert and Fields 2020). This pattern arises because common host variants are selected against by parasite variants that specialize on these host variants. A previous study of the genetic basis of *P. ramosa* infection found usually high levels of genetic diversity at a number of PCL genes, which the authors interpreted as evidence of balancing selection at those loci (Andras et al. 2018). Samples for this previous study were drawn from a wide geographic range that spanned across the Western Palaearctic. Despite the fact that our study samples were drawn from a single population, we observed peaks in nucleotide diversity at the same genomic positions (Fig 2). In respect to the *D. magna* host, we observed the highest levels of nucleotide diversity at the position of the ABC locus, which is known to contain major structural variants of sufficient divergence to limit recombination (Bourgeois et al. 2021). Thus, we observe evidence for balancing selection in terms of nucleotide diversity as well as genomic structural diversity among both the host and parasite dimensions of this system.

### Molecular mechanisms of infection and resistance

Pasteuria collagen-like proteins (PCLs) belong to a larger class of collagen-like proteins which are known to occur in at least 14 bacterial phyla (Qiu et al. 2021). For example, many species of pathogenic *Streptococcus* contain collagen like proteins (Xu et al. 2002; Karlström et al. 2004; Paterson et al. 2008; Lukomski et al. 2017), as does the highly virulent *Bacillus anthracis* (Henriques and Moran 2007). Collagen-like proteins have been directly linked to infectivity in the gut pathogen *Clostridium difficile* (Strong et al. 2014), drug resistant strains of the human and livestock parasites *Burkholderia* sp. (Bachert et al. 2015), and the causative agent of Legionnaire’s disease, *Legionella pneumophila* (Vandersmissen et al. 2010). Collagen-like proteins are hypothesized to mediate the infection process through the mechanism of attachment to host tissues (Qiu et al. 2021). Compared to other bacterial species, *P. ramosa* has an unusually large number of collagen-like proteins, with nearly 50 genes identified thus far (McElroy et al. 2011). The enormous repertoire of collagen-like proteins in *P. ramosa* could be explained as a long-term effect of balancing selection due to a tight coevolutionary history with its host. A noteworthy finding is that the collagen-like genes revealed by our co-GWAS models all appeared in a specific triplet formation. The *P. ramosa* genome contains many isolated collagen genes and loose aggregations of collagen genes, but none of these showed strong association with the host. Thus, with regard to infection phenotype, we observed that not only the nature of the gene is important, but also the spatial orientation of the gene in the genome. The function of the collagen-like gene triplets, as opposed to single genes is not known.

The molecular mechanisms which determine *D. magna* resistance to specific *P. ramosa* genotypes are not yet resolved, although a promising hypothesis concerns the post-translational modification of cuticle proteins via the attachment of branching sugars (*i*.*e*., glycosylation). Previous studies have found major differences between susceptible and resistant clones at the ABC locus in regards to genes encoding fucosyltransferases, glucosyltransferases, galactosyltrasferases – all of which are involved in the post-translational attachment of sugars to proteins (Bento et al. 2017). The E locus, which appears have an independent evolutionary origin from the ABC locus, seems to be enriched for galactosyltransferase and glucosyltransferase genes as well (Ameline et al. 2021). In an interesting parallel, it has been proposed that glycosylation of amino acid residues at the C-terminal domain also plays an important role in attachment specificity of bacterial collagen-like proteins (Strong et al. 2014; Qiu et al. 2021). There is some support for this hypothesis from a previous study of *P. ramosa* infectiousness, which found that a predicted N-glycosylation site in the C-terminal domain of PCL 7 correlated perfectly with infection phenotype (Andras et al. 2020).

### Functional testing of resistance loci

Phenotypic assessment of resistance loci via pasteuria attachment assays has provided important context and support for our co-GWAS model results in several ways. First, it allowed us to test whether any of the uncharacterized peaks in our models corresponded to an identifiable resistance phenotype. The strongest peak recovered across any of our models was localized to the right arm of chromosome 5 in the *D. magna* genome, a position which has not been previously associated with pasteuria resistance. Our functional testing revealed that this genomic region is strongly associated with resistance to pasteuria isolate P20, and thus very likely contains uncharacterized resistance genes. Secondarily, the genomic positions and genetic models for pasteuria resistance were mostly developed through laboratory crosses of *D. magna* from other populations. Functional testing allowed us to verify that pasteuria resistance in a wild population conformed to the patterns expected by our genetic models. Third, functional testing allowed us to determine how many DAPC clusters were necessary to differentiate resistance phenotypes. The imperfect segregation of resistotypes according to our DAPC clusters for the D and E loci show an unexpected departure from the current genetic models of pasteuria resistance, and indicates the possible influence of a yet unknown interacting resistance locus, perhaps one of the uncharacterized loci recovered from our co-GWAS models.

### Generalization of co-GWAS and extension to other systems

The strength of our co-GWAS results are in some part due to the highly specific nature of the interaction between *D. magna* and *P. ramosa*, with specific genotype combinations showing either complete resistance or complete susceptibility to infection. This degree of specificity has allowed us to resolve interlinked host and parasite loci using a relatively modest sample of 258 individuals, which is considerably fewer than typically required to resolve traits of a more continuous nature using conventional GWAS approaches. In general, we expect that the detection power of a co-GWAS is constrained in the same manner as a conventional GWAS; Reasonable detection power will require relatively larger sample sizes in cases where there is low heritability of the trait of interest, complex genomic architectures, or polygenic contributions to a single trait (Korte and Farlow 2013). In the specific context of co-GWAS one should also expect that sample sizes will need to increase as the specificity of the interaction decreases.

Regarding the issue of epistasis in co-GWAS (or GWAS more generally), our study shows that epistatic interactions can be resolved without explicitly providing prior knowledge of epistatic interactions. Our co-GWAS successfully recovered the E-locus for pasteuria resistance, despite the fact that the phenotype encoded by the E-locus is contingent upon the allelic state at the dominant C locus (Fredericksen et al. 2021). While our models did not recover the signal of the A and B loci, which share a similar epistatic relation with the C locus, the underlying issue is more likely a lack of genetic variants rather than insufficiency of the model itself as these loci appear to be nearly fixed in our study population (Ameline et al. 2021).

In conclusion, understanding the genomic basis of infectious disease is a fundamental objective in the study of coevolution. Our results show a clear signal of interspecies linkage disequilibrium across multiple sets of interacting loci in a naturally coevolving system, thus confirming the most fundamental assumption of host-parasite coevolution. Previous co-genomic association studies have identified variants of clinical significance for HIV infections (Bartha et al. 2013), Hepatitis C (Ansari et al. 2017), pneumococcal meningitis (*16*), and malaria (Band et al. 2021) through a combination of SNP arrays and patient clinical data. Our approach moves a step further by examining the complete genomes of naturally infected hosts and parasites from a host population with a known history of parasite epidemics. Our central finding is an underlying host-parasite genetic interaction matrix, as such matrices are at the core of coevolutionary theory, and their topology determines the tempo and mode of coevolution. Our matrix confirms the importance of multiple loci in both antagonists, acting together within and across genomes, with epistasis playing a decisive role for the host and the parasite. We demonstrate that the co-genomic framework can be successfully applied to non-model systems, and given sufficiently strong host-parasite specificity, resolution of complex traits is possible in a candidate-free manner. In the future, this method will be useful to elucidate other types of pairwise interactions, such as commensalism, mutualism, predation, and assortative mating in the context of sexual selection.

## MATERIALS AND METHODS

### Field Collections

We collected *D. magna* samples from Lake Aegelsee (47.558048, 8.861063) via horizontal net tows at the deepest place during a strong *P. ramosa* epidemic in summer 2019. These wild-collected daphnia were monitored in the laboratory for signs of *P. ramosa* infection and the appearance of offspring. Although *P. ramosa* typically castrates the host, clonal reproduction is still possible in the early stages of infection. As *P. ramosa* is exclusively transmitted through the environment, the laboratory lines were quickly separated from the mother and remained uninfected throughout the experiment. After verifying *P. ramosa* infection status and establishing healthy clonal lines, we preserved the original wild-collected individuals in 90 % EtOH at -20 °C for later DNA extraction.

### DNA extraction and Sequencing

Whole-organism DNA extractions from daphnia may contain a high proportion of non-target DNA from undigested food, gut microbiota, and bacteria microfilm on the cuticle. To reduce contaminants, we cleared the gut by feeding the daphnia with a suspension of sterile beads (Sephadex G-25, cross-linked dextran gel) three times per day for at least 24 hours. After the gut was visibly cleared of food, we monitored containers with individual daphnia every 2-4 hours for the presence of a molted cuticle. After molting, daphnia were immediately preserved in 90 % EtOH at -20 °C for later DNA extraction, sharply reducing the amount of bacterial contamination in the preserved sample. Preliminary experiments showed this protocol to be effective in limiting the amount of non-target DNA in preserved samples to less than 5 %.

We extracted DNA from infected daphnia using a modified protocol for a Gentra PureGene column-based extraction kit. Briefly, we homogenized whole daphnia with a pestle and incubated the samples overnight at 55 °C in a solution of lysis buffer and Proteinase K. The samples were then incubated for 30 minutes at 37 °C with RNase A, treated with a protein precipitation solution, and centrifuged to remove precipitates. The supernatant was then subject to a chilled isopropanol precipitation with glycogen added as a DNA carrier. The DNA pellet was then resuspended in a buffer solution and stored at -20 °C. Our complete extraction protocol is available on the GitHub repository for this manuscript (https://github.com/edexter/Interlink).

We then prepared genomic libraries for short-read sequencing using a NEBNext Ultra II DNA Library Prep Kit for Illumina. The libraries were sequenced across multiple runs of a Novaseq 6000 to generate paired-end 150 bp reads, aiming for at least 30X coverage across host and parasite genomes. Samples were sequenced with individual barcodes for subsequent demultiplexing and inspection for sequencer batch and lane effects. Genomic library preparation and Illumina sequencing were performed at the Basel Genomics Facility, Basel, Switzerland.

### Host phenotyping

The most important step of the *P. ramosa / D. magna* infection process is attachment of *P. ramosa* spores to the any of several sites on the cuticle, and may occur through the uptake of *P. ramosa* spores during feeding (Duneau et al. 2011; Fredericksen et al. 2021). Attachment is a necessary requirement for successful infection and variation in attachment explains the vast majority of overall infection success (Hall et al. 2019, Molecular Ecology). Attachment specificity between host and parasite genotype combinations can be determined via a relatively simple attachment test using fluorescently labeled pasteuria spores. Experimental studies based upon this attachment test have found a nearly perfect correlation between attachment and infection outcome, with little to no effect of environmental or host condition (Duneau et al. 2011).

We collected resistance phenotype data of the host (aka. “resistotype”) for a subset of 107 genotypes of our sequenced *D. magna* clones, with each clone tested in triplicate. Daphnia resistotype was assessed through gut attachment tests with fluorescently labeled pasteuria spores using a standardized panel of five pasteuria isolates: C1, C19, P15, P20, and P21 (Duneau et al. 2011; Bento et al. 2017; Bento et al. 2020; Ameline et al. 2021). Resistance (R) or susceptibility (S) to each isolate is denoted in a five-letter resistotype code with each position corresponding to a single isolate. For example, the resistotype [SRRRR] would indicate susceptibility only to C1, while the resistotype [SSSRR] would indicate susceptibility to C1, C19, and P15. These attachment data were used for *post hoc* validation of our co-GWAS results through comparison of genotypes at interlinked loci and resistance phenotypes.

### Bioinformatics

De-multiplexed sequence-read pairs were trimmed to remove low-quality reads and adaptor sequences using Trimmomatic version 0.39 (Bolger et al. 2014). Duplicate reads were then removed using Picard version 2.9.2. The trimmed reads were mapped against host and parasite reference genomes using BWA-MEM version 0.7.15 operating under default parameters (Li 2013). Reads were mapped to the daphnia XINB version 3.0 draft genome assembly and the pasteuria isolate C1 version 1.0 draft genome assembly (manuscripts in prep. – available by request). Reads mapping to both host and parasite genome were flagged for downstream filtering. We jointly called genomic variants (SNPs, indels, and mixed-type variants) across all samples using the GATK haplotype caller following the best practices pipeline for diploid samples (Auwera and O’Connor 2020). Although *P. ramosa* is haploid, this approach enabled us to differentiate single genotype infections from multiple infections, and to identify major and minor alleles within each pool. *In silico* experiments showed little change in the major/minor alleles identified when calling *P. ramosa* variants under different ploidy models.

The host and parasite VCF files were filtered to remove low-quality bases, poor alignment scores, or extreme strand biases. Variants were then filtered to exclude sites with excessive depth (> 2x mean depth), or excessive instances of dual mapping (reads mapping to both genomes) using a custom R script. Based upon visual inspection of dual mapping regions, low-complexity sequences rather than assembly errors or gene homology appear to be the most likely cause of dual mapping. The VCF files were then filtered to remove multi-allelic variants and converted to PED files for input to Plink for co-GWAS analysis. Summary statistics related to read mapping were calculated with SAMtools version 1.10 (Li et al. 2009). We calculated nucleotide diversity (π) for host and parasite genomes with VCFtools version 0.1.16 (Danecek et al. 2011). For the host, we calculated π across 100 Kb sliding windows with a 1 Kb offset. Across the much smaller parasite genome, we calculated π across 500 bp sliding windows with a 250 bp offset. Precise filter cutoff values and other parameters can be found along with our complete code at the associated Github repository.

### Co-GWAS analysis

We employed a co-genome wide sequencing association study (co-GWAS) framework to search for interlinked genomic regions of the host and parasite. This approach is based on the regression of host genotypes against parasite genotypes for every pairwise combination of loci. Similar methods of genome-to-genome association testing have been previously employed to find clinically relevant variants in HIV (Bartha et al. 2013), Hepatitis C (Ansari et al. 2017), *Streptococcus pneumoniae* (Lees et al. 2019), and *Plasmodium falciparum* (Band et al. 2021) using human SNP array data. We prepared our genomic data for co-GWAS by removing variants with a minor allele frequency less than 0.05 or missingness greater than 0.20. Pairwise relatedness was calculated for the host using the KING kinship estimator, and samples showing relatedness greater than 0.35 (possible clone mates) were thinned to remove redundant samples. To reduce the probability of false positives due to population structure, we conducted PCA on subsets of linkage disequilibrium thinned loci for both host and parasite. Loci were thinned using the indep-pairwise function in Plink with a maximum r2 threshold of 0.2 across a 1 kb (pasteuria) or 1000 kb (daphnia) sliding window. A smaller window size was selected for the parasite due to its relatively small genome. The top principal components were then included as covariates in our co-GWAS models. We used Plink version 2.00a3LM for PCA, frequency-based filtering, and co-GWAS analysis (Chang et al. 2015).

Our GWAS analysis was structured as a logistic regression model of the form:

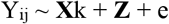

Where *Y* is a binary variable [0,1] indicating the presence or absence of allele *i* at locus *j* in the parasite genome. For example, the biallelic SNP A/T would be split into 2 variables. A pasteuria sample containing only at the A allele would be coded as 1 for the first variable and 0 for the second, a pasteuria sample showing only the T allele would be coded as 0 and 1 respectively, and a multiple infected sample showing both alleles (i.e. a heterozygote genotype call) would be coded as 1 and 1. The variable *X* represents the genotype at locus *k* in the host genome expressed as the major allele dosage [0,1,2], a dominant reference allele model [1,1,0], or a dominant alternate allele model [0,1,1]. These different coding schemes reflect the fact that host samples contain a single diploid individual while their infecting parasites may contain a pool of haploid individuals (i.e., multiple infections). Note that although dominance models are more biological plausible than a simple genotypic dosage model, they are rarely employed in the context of GWAS due to statistical difficulties in detecting non-additivity (i.e., too few homozygous-minor genotypes) (Chang et al. 2015; Marees et al. 2018). The term *Z* [0-1] represents a matrix of the top *n* principal components and controls for population structure among samples, with permutations of our primary co-GWAS model containing different subsets of principal components. In order to reduce the computational burden of weak or non-significant associations, test results with p-values greater than 10^−4^ were not stored for downstream analysis or plotting. This cutoff was derived from preliminary co-GWAS trials which showed that p-values below this threshold were uniformly distributed across the daphnia genome and generally uninformative.

Because each variant position in the host genome is tested against each variant position in the parasite genome, co-GWAS analysis can generate a considerable number of false positives purely through stochastic processes. Rather than applying a simple threshold to denote statistical significance, we utilized information about the magnitude and breadth of each peak to create a list of candidate loci for further examination. Specifically, biologically relevant associations are expected to yield a broad peak of elevated p-values, including nearby flanking regions which rapidly decay with distance from the site of selection. This pattern is a characteristic signature of linkage disequilibrium decay due to recombination and provides a useful biological validation to our statistical results. In contrast, we expect that associations due to mapping errors or stochastic sampling effects will result in isolated peaks comprised by one or very few variants.

### Host and parasite genotype clustering

We extracted the sequence data from the genomic region corresponding to each of the known host resistance loci and applied a discriminant analysis of principal components (DAPC) function to aggregate individual genotypes into clusters of broadly similar genotypes (Jombart et al. 2010). Genotype clusters were then grouped by resistance phenotype to define functional groups. Each host was assigned a set of three resistance locus codes denoting the functional group of the C, D, and E locus, each with two functional groups (e.g., C vs C’). We applied the same DAPC method to our parasite samples, using the entire parasite genome rather than individual loci. DAPC and PCA were performed using the Adegenet version 2.1.3 package for R version 4.0.4 (Jombart 2008). All statistical analyses were performed using R unless specifically mentioned otherwise, and all figures were produced using the ggplot2 (version 3.3.3) package (R Core Team 2016; Wickham et al. 2018). DAPC clustering results were compared against genomic clusters generated via the K-means cross-validation algorithm implemented in Admixture version 1.3.0 (Alexander et al. 2009). Admixture analysis was performed using a set of LD-thinned biallelic variants from singly-infected samples, with cross validation set to select the optimal value for K ranging from 1 to 10.

## Acknowledgments

We would like to thank the Ebert lab members for valuable discussions and suggestions.

## Author contributions

Conceptualization: ED, DE, PDF

Methodology: ED, DE, PDF

Investigation: ED, DE

Visualization: ED

Funding acquisition: ED, DE

Project administration: ED, DE

Supervision: DE

Writing – original draft: ED, DE

Writing – review & editing: ED, DE, PDF

## Data and materials availability

The complete annotated code necessary to reproduce all analyses and figures presented here is available on GitHub (https://github.com/edexter/Interlink). The raw genomic data pertaining to this study are available from the NCBI Short Read Archive (SRA) (accession number in progress).

