## Supplemental tables and figures for "Uncovering the genomic basis of infection through co-genomic sequencing of hosts and parasites"

**Affiliations:**

**S1. (A)** The percentage of sequencer reads mapping to either the *D. magna* host or *P. ramosa* parasite genome. Each sample corresponds to a single *D. magna* naturally infected by *P. ramosa*. Unmapped reads likely represent gut content or microbiota. **(B)** Scatterplot of coverage across host and parasite genomes. Each point represents a single sample. The y-axis shows the proportion of the reference genome covered by sequencer reads from a single sample, with the red line indicating a 95 % cutoff for inclusion in the final data set. A total of 10 samples were excluded from further analysis based on this criterion. 8 of these samples failed to reach the coverage threshold due to low recovery of pasteuria DNA, while the remaining 2 samples were excluded due to technical issues with library preparation.

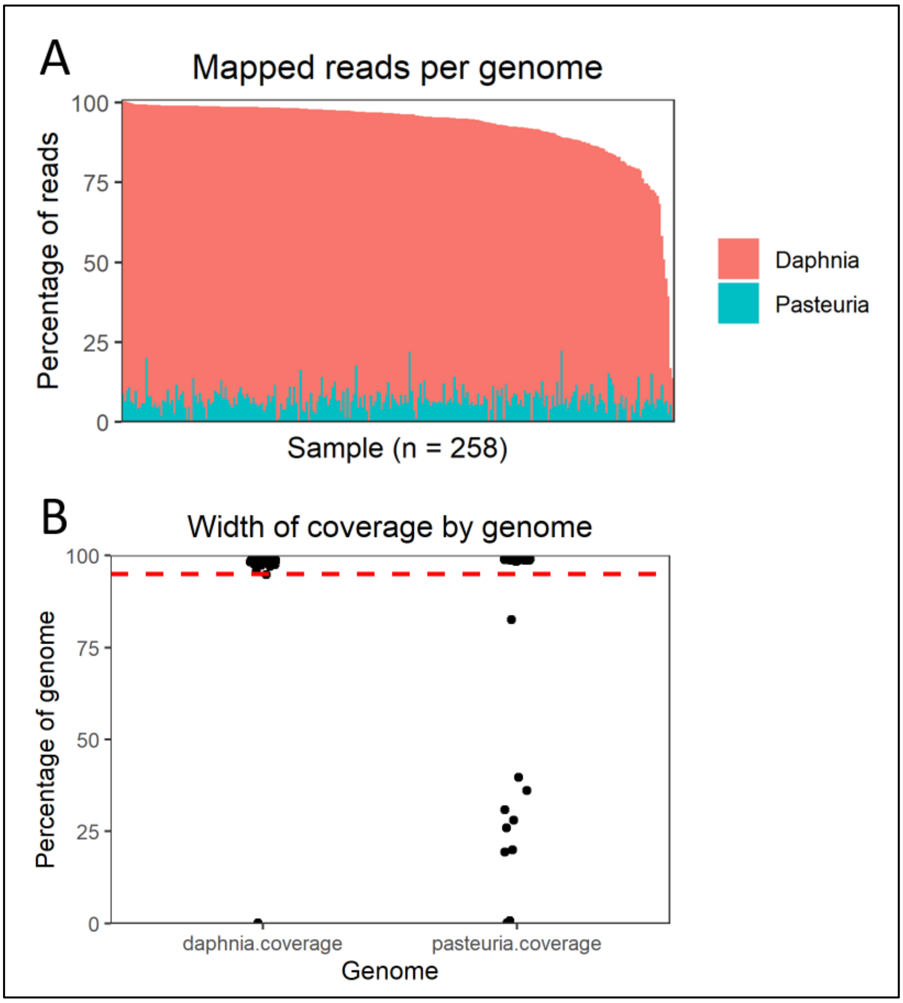

**S2.** Admixture derived clusters of *P. ramosa* samples, excluding clear cases of multiple infection. Multiple infections were defined based on elevated within-sample polymorphism (see main text). A) Model cross-validation shows that the majority of variation can be explained by three groups. B) PCA ordination of the three admixture groups, which corresponds exactly to the *Alpha*, *Beta*, and *Gamma* lineages identified through DAPC clustering. C) Admixture scenarios with greater than 3 groups support gene flow within the three DAPC lineages, but not across them. Here K = 7 is shown for example. Note that PC 1 and PC 3 are shown in both sub-panels in order to visualize *Alpha* substructure.

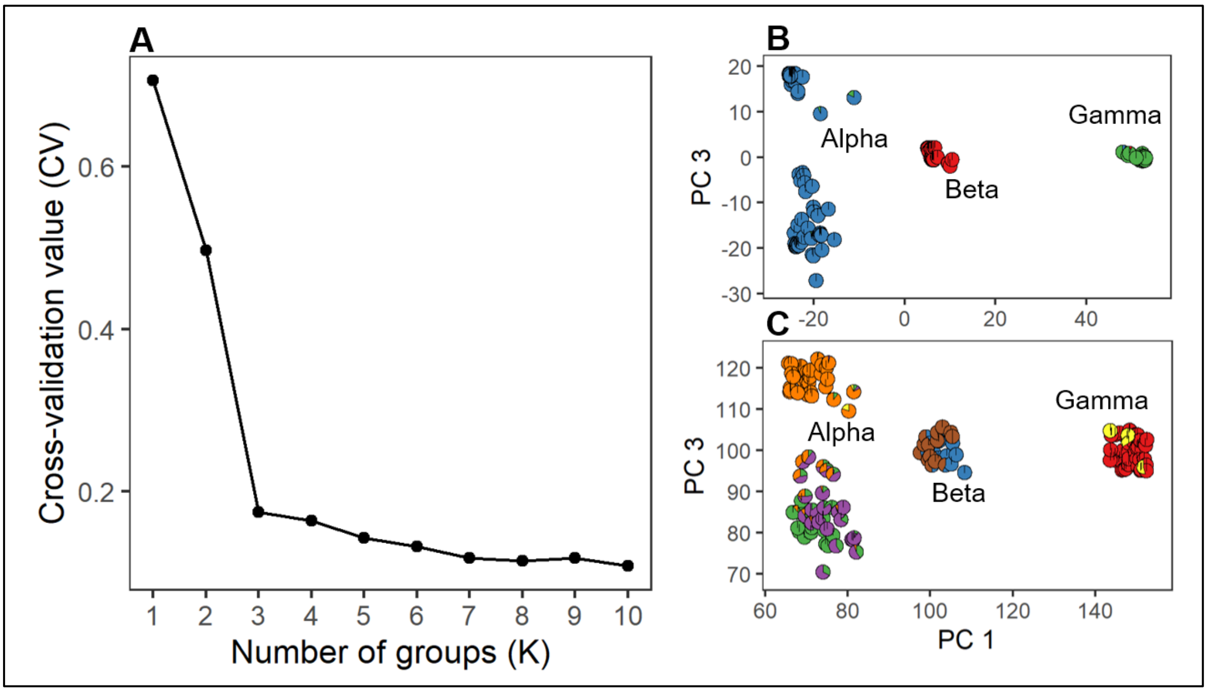

**S3.** Plot of linkage disequilibrium dissolution with distance across the *P. ramosa* genome. Pairwise LD was calculated for all variants across the genome and then mean values were calculated for 100 bp bins. Results are shown for the C1 reference genome and an additional reference genome generated from the distantly related *P. ramosa* isolate P21. LD decay is strongly observed across relatively short distance, which is an indicator of ongoing genetic recombination. Note that LD values do not reach zero in this dataset as some of the samples in this dataset are likely clonemates.

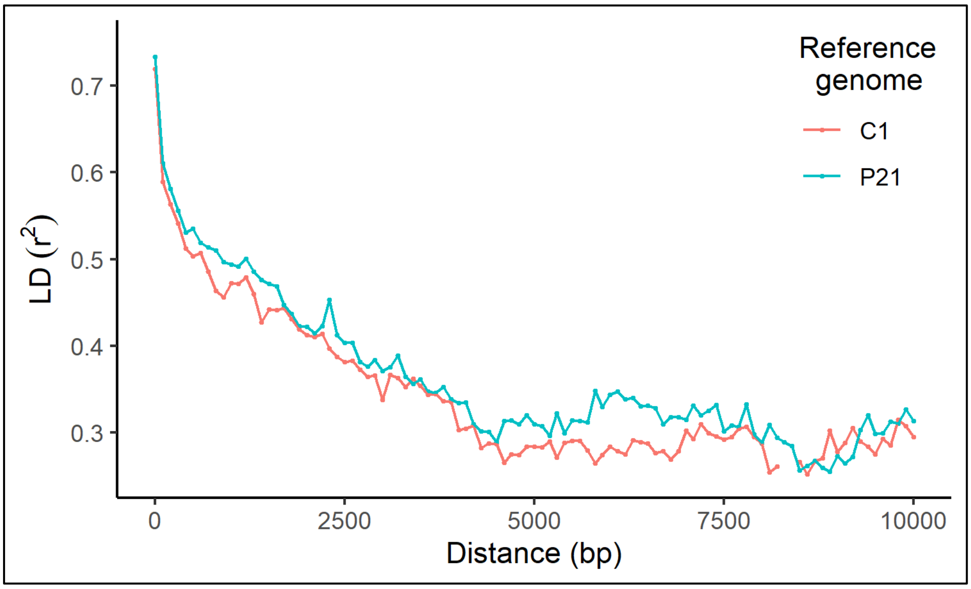

**S4.** Pasteuria Collagen-Like (PCL) gene positions in the genome assembly of *P. ramosa* strain C1. Complete PCL triplets comprised by a single member from each of the main PCL subfamilies are highlighted in blue. Alternate shades separate adjacent triplets on the table. Due to the circular nature of the *P. ramosa* genome, PCLs located at the bottom of the table are located in relatively close proximity to PCLs at the top of the table. Note that PCL subfamily 4 may have a different evolutionary origin than the other subfamilies.

| Start position | End position | Name | PCL subfamily |
|---|---|---|---|
| 7278 | 8312 | Pcl6 | 1 |
| 8355 | 9347 | Pcl7 | 2 |
| 9393 | 10679 | Pcl8 | 3 |
| 59538 | 60728 | Pcl34 | 4 |
| 60761 | 61723 | Pcl35 | 4 |
| 61764 | 61886 | Pcl8 | 3 |
| 62275 | 63426 | Pcl3 | 3 |
| 86443 | 87708 | Pcl23 | 3 |
| 87759 | 88736 | Pcl22 | 2 |
| 89000 | 90028 | Pcl21A | 1 |
| 138861 | 139442 | Pcl33 | 4 |
| 241583 | 242773 | Pcl5 | 4 |
| 242861 | 243832 | Pcl4 | 2 |
| 243877 | 245130 | Pcl3 | 1 |
| 317101 | 317544 | Pcl21B | 1 |
| 317594 | 318541 | Pcl27 | 2 |
| 318592 | 319830 | Pcl28 | 3 |
| 380105 | 380875 | Pcl32 | 4 |
| 775391 | 776653 | PclBac | ? |
| 780508 | 781203 | Pcl31 | 4 |
| 874337 | 875257 | PclBac2 | ? |
| 1028703 | 1030220 | Pcl1a | 4 |
| 1030535 | 1031569 | Pcl2 | 4 |
| 1199952 | 1201148 | Pcl15 | 1 |
| 1201186 | 1202148 | Pcl16 | 2 |
| 1202192 | 1203511 | Pcl17 | 3 |
| 1325522 | 1326673 | Pcl29 | 1 |
| 1478817 | 1479578 | Pcl20 | 4 |
| 1480175 | 1480939 | Pcl19 | 4 |
| 1481032 | 1481940 | Pcl18 | 4 |
| 1483210 | 1483965 | Pcl30 | 4 |
| 1509135 | 1510478 | PclBac3 | ? |
| 1517919 | 1519055 | Pcl24 | 1 |
| 1519111 | 1520061 | Pcl25 | 2 |
| 1520134 | 1521372 | Pcl26 | 3 |
| 1571213 | 1572475 | Pcl9 | 1 |
| 1572528 | 1573523 | Pcl10 | 2 |
| 1573552 | 1574559 | Pcl11 | 3 |
| 1577719 | 1578984 | Pcl12 | 3 |
| 1579022 | 1579993 | Pcl13 | 2 |
| 1580131 | 1581168 | Pcl14 | 1 |
| 1586627 | 1587892 | Pcl38 | 3 |
| 1587930 | 1588910 | Pcl37 | 2 |
| 1588930 | 1589937 | Pcl36 | 1 |

**S5.** Detailed view of Manhattan plot peaks for co-GWAS model 1. Panel A shows a broad peak that spans > 100 Kb with a clear dissolution of LD with distance. This is the expected signature of a true biological peak when performing co-GWAS with un-thinned whole genome data. The coordinates of the ABC locus region are shown in blue. The well-characterized non-homologous region of the ABC locus is shown by a gap in the middle of the locus where few reads are mapped. Haplotypes at this position are so divergent that mapping produces an apparent gap in the middle of the locus, with no statistics being computable. Panel B shows a contrasting, and far less convincing pattern, with the peak represented by one apparently intragenic SNP absent of linked variation. This peak likely represents a mapping artifact or stochastic noise in the dataset.

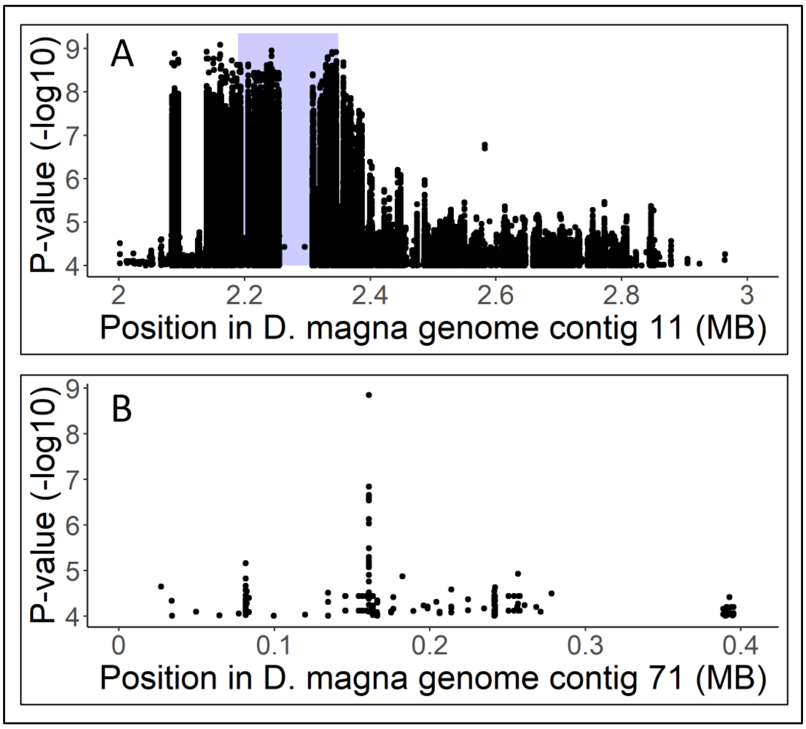

**S6.** Co-GWAS model 3 results filtered to show the top 100 associations for contigs 11 and 18 of the *D. magna* genome, which respectively contain the C and D loci for pasteuria resistance. Although these loci exhibit similar pattern of associations as shown in models 1 and 2 (Figure 2 main text), the signal is relatively weak and difficult to discern in model 3 compared to the strong peak of association arising from chromosome 5.

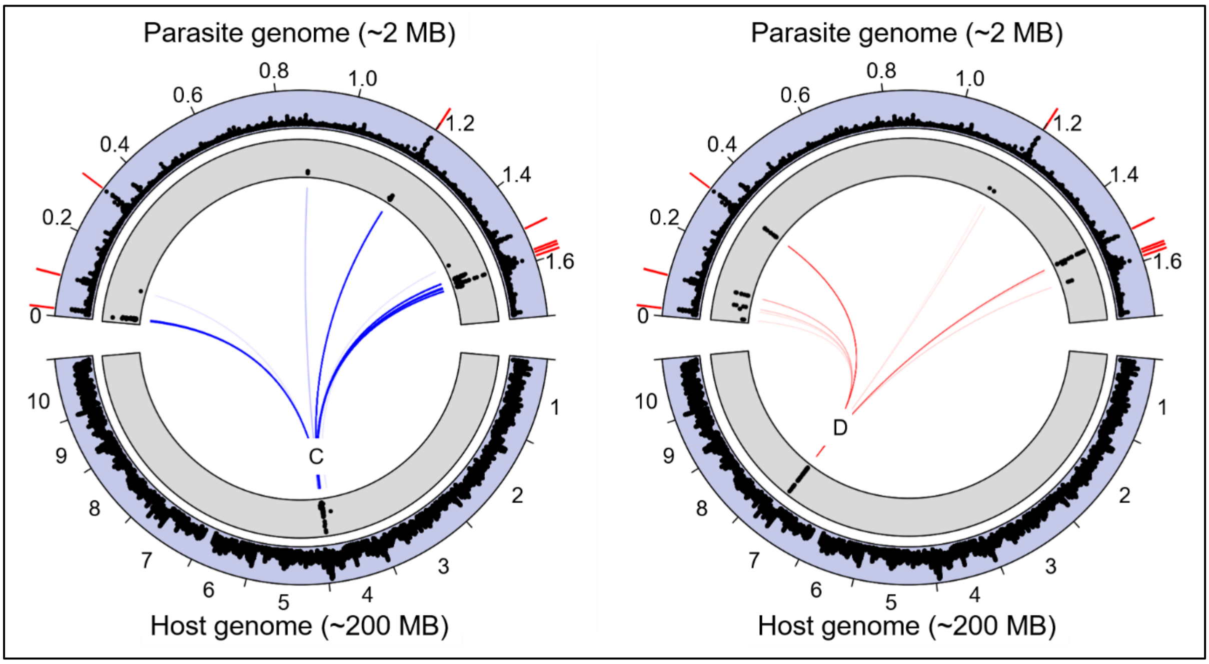

**S7.** Genotype clusters from DAPC analysis of all variant positions located within the ABC locus. Resistotypes (2nd and 3rd columns) are shown relative to the C1 and C19 *P. ramosa* isolates. Only samples with RR and SS resistotypes (94 % of total) are shown. The minimum number of groups required to segregate resistotypes was 3, perhaps reflecting the two homozygote and the heterozygote genotypes at these loci.

| ABC locus genotype cluster | C1/C19 RR | C1/C19 SS | Genotype group shown on figure 4 | Possible allelic state according to Bento 2017 |
|---|---|---|---|---|
| Group 1 | 57 | 0 | C | CC, Cc |
| Group 2 | 19 | 0 | C | CC, Cc |
| Group 3 | 0 | 24 | C' | cc |

**S8.** Genotype clusters from DAPC analysis of all variant positions located within the D locus. Resistotypes (2nd and 3rd column) are shown relative to the P15 and P21 *P. ramosa* isolates. Note that the low number of P21 resistant hosts makes determination of the allelic state for genotype 2 somewhat tentative.

| D locus genotype cluster | P15/P21 RR | P15/P21 RS | P15/P21 SR | P15/P21 SS | Genotype group shown on figure 4 | Possible allelic state according to Bento 2020 |
|---|---|---|---|---|---|---|
| Cluster 1 | 0 | 2 | 2 | 50 | D | DD, Dd |
| Cluster 3 | 0 | 1 | 2 | 20 | D | DD, Dd |
| Cluster 2 | 4 | 5 | 2 | 18 | D' | DD, Dd, dd |

**S9.** Genotype clusters from DAPC analysis of all variant positions located within the E locus. Resistotypes (2nd and 3rd column) are shown relative to the C1, C19, and P20 *P. ramosa* isolates. Only C1/C19 RR resistotypes are shown, i.e. homozygotes or heterozygotes with the dominant C allele at the C locus, because homozygosity of the alternative c-allele nullifies the effect of the E-locus due to epistasis (Ameline et al. 2021).

| E locus genotype cluster | C1/C19/P20 RRR | C1/C19/P20 RRS | Genotype group shown on figure 4 | Possible allelic state according to Ameline et al. 2021 (13) |
|---|---|---|---|---|
| Cluster 2 | 38 | 0 | E | [CC, Cc] [ee] |
| Cluster 3 | 16 | 0 | E | [CC, Cc] [ee] |
| Cluster 1 | 9 | 13 | E' | [CC, Cc] [EE, Ee, ee] |

**S10.** DAPC genomic clusters based on the sequence data at each known resistance locus region in the *D. magna* genome. Up to 10 genotype clusters could be discerned at each resistance locus, likely representing several haplotypes in various combinations with each other. In all cases, we found that the maximum segregation of resistotypes could be achieved with three clusters, which could be further collapsed into two groups representing primarily resistant or primarily susceptible phenotypes. Imperfect correspondences between genotype and phenotype at the D and E loci do not appear to be an artifact of our clustering methodology, and may reflect the influence of yet unknown interacting resistance loci.

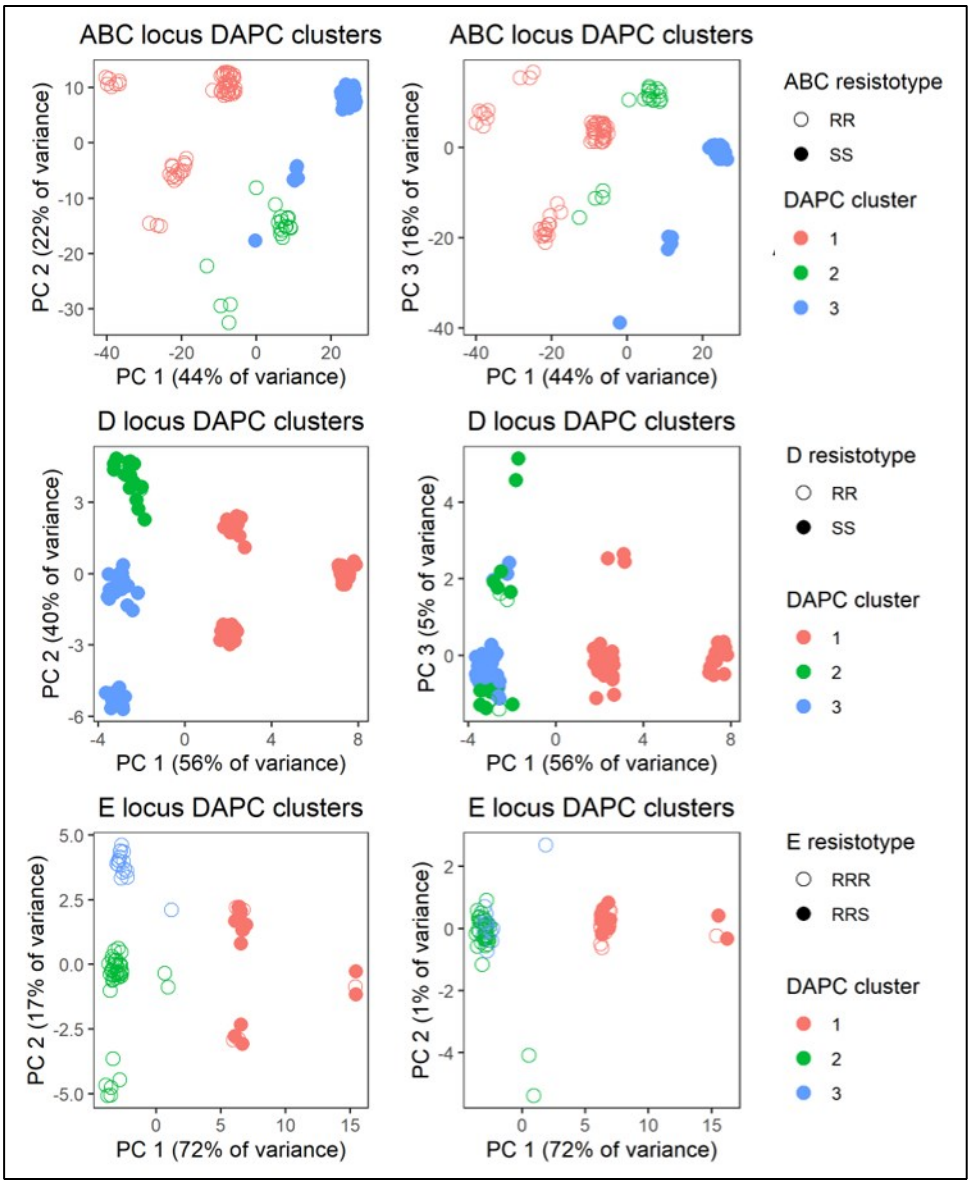

**S11.** PCA loading score density across the *D. magna* genome relative to principal component 2. The vertical lines represent the location of the C, E, and D loci from left to right. The dashed line shows the density function adjusted to correct for the distribution of variants across the genome. PC 2 is strongly driven by local variation at the position of the E locus.

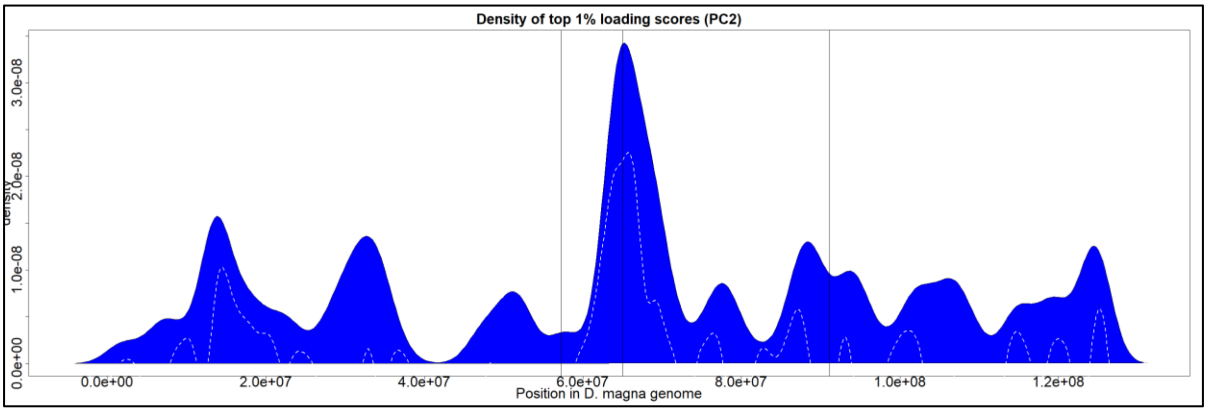

**S12.** Co-GWAS "Megalopolis" style plots for host and parasite genomes. The co-GWAS framework can be considered as a series of single GWAS models that treat the presence or absence of each variant in the paired genome as a distinct "phenotype". These results can be visualized as a series of stacked Manhattan plots which can be viewed simultaneously as if one were looking down through a very large stack of transparent pages ($10^4$ – $10^5$ layers for each panel). This approach can be instructive as an exploratory data analysis tool for co-GWAS, but can be misleading when interpreted as a typical Manhattan plot; hence our preference for the slightly different nomenclature. Here we see the correspondence between the raw co-GWAS data visualized in this Megalopolis plot format, and the circular plot format shown in the main text which more explicitly shows the strongest points of interlinkage between the genomes.

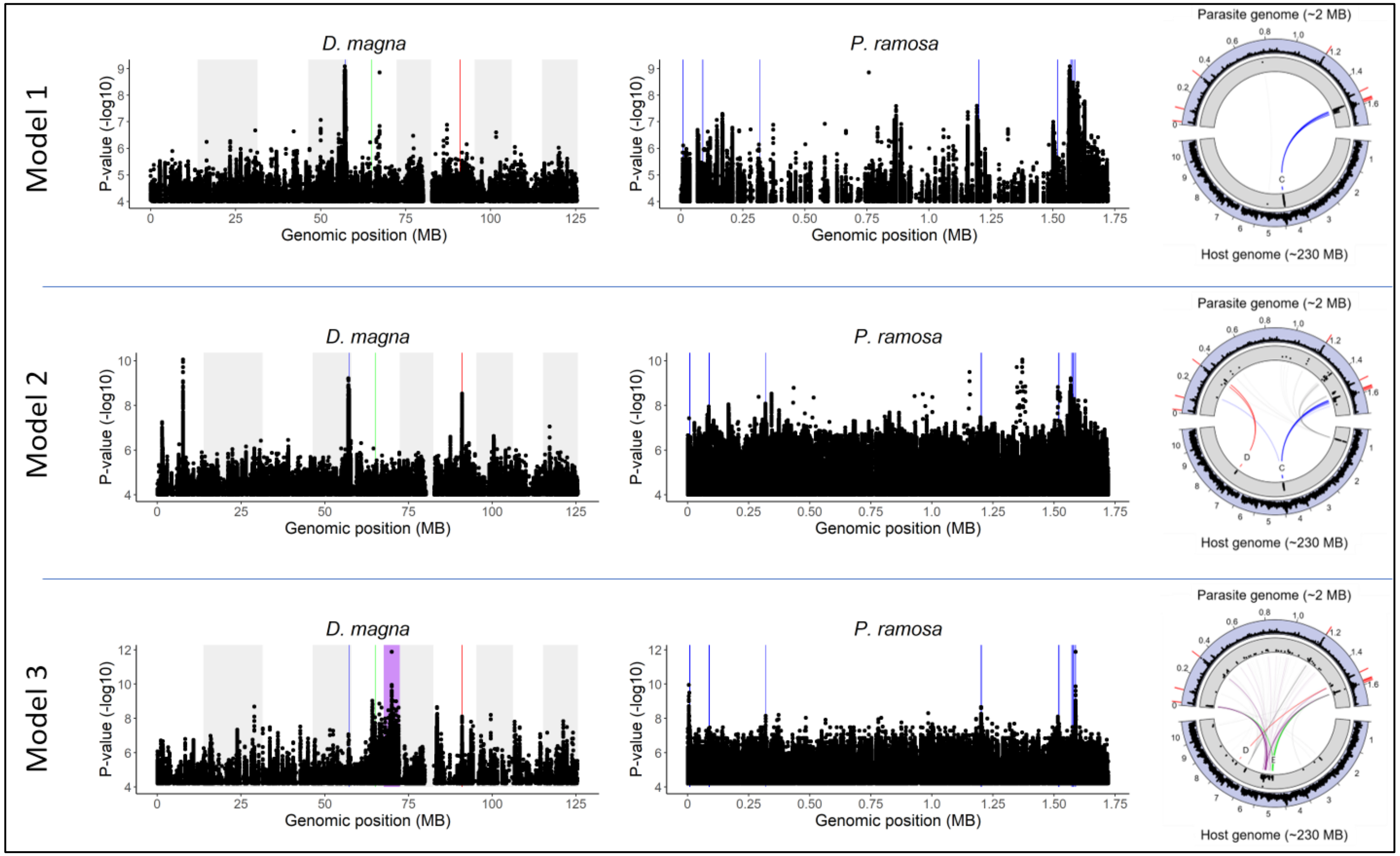
